# DC-SIGN Binding to the Surface Ig Oligomannose-type Glycans Promotes Follicular Lymphoma Cell Adhesion and Survival

**DOI:** 10.1101/2025.07.11.664379

**Authors:** Giorgia Chiodin, Dylan Tatterton, Philip J. Rock, Luis del Rio, Erin Snook, Sonya James, Patrick J. Duriez, Miriam Di Re, Martijn Verdoes, Stuart Lanham, Daniel J. Hodson, Richard W. Burack, Francesco Forconi

## Abstract

The occupation of the surface immunoglobulin antigen-binding site by oligomannose-type glycans (sIg-Mann) is a tumor-specific post-translational modification of classic follicular lymphoma (FL). SIg-Mann switches binding from antigen to dendritic cell-specific intercellular adhesion molecule 3 grabbing non-integrin (DC-SIGN), known to be expressed on interfollicular macrophages and FL-associated follicular dendritic cells (FDCs). The interaction with DC-SIGN induces reorganization of sIg-Mann in wider and less dense clusters than anti-Ig, consistent with inefficient DC-SIGN-induced endocytosis and a low-level intracellular signaling. However, ligand-specific cell clusters form between sIg-Mann-expressing lymphoma and DC-SIGN-expressing cells, raising a need to understand the functional consequences of the interaction of DC-SIGN with sIg-Mann on primary FL cells. This engagement induces adhesion of FL cells to VCAM-1 *via* B-cell receptor proximal kinases and actin regulators in a fashion similar to anti-Ig, but without initiating apoptosis *in vitro*. Instead, antibody blockade of sIg-Mann contact with DC-SIGN expressed on FDC-derived YK6/SIGN cells inhibits adhesion and survival of primary FL cells *in vitro*. These data highlight that the specific interaction with DC-SIGN induces FL cell adhesion to VCAM-1, likely allowing FL cell retention in the lymph node, and survival of the FL cells. Adhesion and survival are inhibited by an anti-DC-SIGN blocking antibody, indicating a new early therapeutic approach against FL retention and survival in adaptive tumor tissue niches.

**Key points:**

1. DC-SIGN promotes FL cell adhesion to VCAM-1 *via* B-cell receptor proximal kinases and actin regulators
2. DC-SIGN interaction with sIg-Mann sustains the survival of FL cells

## INTRODUCTION

The surface immunoglobulin (sIg) is the fundamental receptor defining the fate of normal and mature tumor B cells.^1^ In classic follicular lymphoma (FL), the sIg undergoes a mandatory tumor-specific post-translational modification. This is the occupation of the antigen-binding site by oligomannose-type glycans.^2^ This unique sIg structure containing oligomannose-type glycans (sIg-Mann) is not detected in circulating and tissue non-tumor B cell subsets,^3^ and results from the acquisition of N-glycosylation sites at specific locations of the complementarity-determining regions (CDR) by somatic hypermutation.^3-5^ The sites are required in the entire clonal population, and if one site is lost, which is rare, either the progeny disappears or is rescued by a new site.^3,6^ The sites are observed from the early stages of FL *in situ* through transformation into DLBCL, where the presence of sIg-Mann is an independent marker of rapid progression.^3,5,7^

The location of the oligomannose-type glycans at the CDR is a positive selection process critical to determining two important changes. The first is to prevent sIg binding to conventional antigens.^8^ In reactive follicles, antigen promotes sIg endocytosis and, in the absence of T-cell help and a rigorous selection by follicular dendritic cells (FDCs), B cell apoptosis.^9-11^ The second critical change is to recognize dendritic cell-specific ICAM-3–grabbing nonintegrin (DC-SIGN).^3,12-15^ This lectin is known to be expressed on interfollicular macrophages and, as studies in FL have shown, follicular dendritic cells (FDCs).^2,16-18^ In contrast to reactive follicles, the sIg levels of FL cells are high while somatic hypermutation is ongoing, and there is little evidence of endocytosis or cell death.^19^ DC-SIGN does not promote sIg endocytosis *in vitro*,^13,14^ and induces a prolonged low-level antigen-independent signal, skewed towards AKT and MYC,^14,15^ resembling the constitutive signaling essential for B-cell survival.^1,20^ This type of signaling is accompanied by a reduced redistribution of sIg on the cell membrane.^21^ However, the consequences of DC-SIGN interaction with sIg-Mann on membrane organization and FL cell fate have not yet been tested.

FL is typically characterized by the retention of the tumor B cells in the lymph node, where they survive within a reticulum of cells, including the DC-SIGN-expressing macrophages and FDCs.^7,16,22,23^ Lymphoma cells expressing sIg-Mann cluster around DC-SIGN-expressing cells.^3^ Antibodies blocking DC-SIGN at the carbohydrate recognition domain inhibit or interrupt the clusters, suggesting a specific role of DC-SIGN in tissue retention.^3^ However, the contribution of the DC-SIGN interaction with sIg-Mann on FL cell retention and survival is not known.

This study explored the role of DC-SIGN in regulating sIg-Mann membrane distribution, FL cell adhesion, and survival. Our findings demonstrate that DC-SIGN significantly enhances adhesion to VCAM-1, expressed by the lymph node reticulum, and provides a survival advantage to FL cells, which is lost when the interaction between DC-SIGN and FL oligomannose-type glycans is blocked.

## METHODS

### Primary samples, cell lines, phenotypic, immunoglobulin gene and intracellular signaling analyses

Details about primary samples, cell lines, phenotypic studies including recombinant DC-SIGN binding and intracellular signaling analyses, and *IGHV-IGHD-IGHJ* transcript analysis are described in the **Supplemental Methods**. The study was approved by the University of Southampton (H228/02/t) and the University of Rochester (RSRB#159) Institutional Review Boards. All patients provided informed consent.

### Direct stochastic optical reconstruction microscopy (dSTORM)

WSU-FSCCL cells (1×10^6^) were incubated in 100 µl calcium-enriched medium (RPMI with 10% FBS and 1 mM calcium) with 20 µg/ml soluble DC-SIGN-Fc or soluble polyclonal goat F(ab’)^2^ anti-human Igκ (Southern Biotech) in 96-well U-bottom plates on ice for 30 minutes, and at 37°C for 10 minutes. Cells were transferred into FACS tubes with ice-cold PBS and fixed with 4% paraformaldehyde (PFA) on ice for 20 minutes. Cells were washed in PBS and stained with 50 µg/ml AF647-conjugated Fab anti-human IgM (Jackson Immunoresearch) for 1 hour at room temperature. Cells were washed in PBS/Tween and PBS, and transferred to Ibidi 8-well glass bottom chambers (Thistle Scientific) which had been sequentially washed with 70% ethanol, acetone and water, coated with 0.01% poly-D-lysine (Sigma Aldrich), and centrifuged at 250G for 10 minutes.

PBS was removed from the well, TCEP STORM buffer^24^ was added, and dSTORM images were acquired. Images (10,000 frames, 30 milliseconds exposure) were collected with an illumination angle of 52° (TIRF) using the ONI Nanoimager equipped with a 640 nm laser and NimOS1.18.3 software (ONI, UK), and analyzed using the CODI cloud analysis platform (beta version, ONI, UK). Images were subjected to drift correction and filtering before regions of interest (ROI) were drawn around the cells. Localizations represent the positions of detected fluorophores, which are identified by the software by fitting high-intensity photon spots to a Gaussian function. Localizations within the ROI were identified and grouped into clusters using HDBSCAN (https://hdbscan.readthedocs.io/en/latest/). The following features were extracted for each cluster: number of localizations; area (computed from the convex hull of the cluster); and density (localizations/area).

### Adhesion assay

Adhesion assays were performed in 96-well (flat bottom) plates (Corning) coated overnight at 4°C with 10µg/ml recombinant human VCAM-1 (Peprotech) in PBS. Plates were blocked for 1 hour in RPMI + 4% bovine serum albumin (BSA) before the experiment. Cells were resuspended in calcium-enriched medium (RPMI with 1% FBS and 1 mM calcium) and incubated with 20 µg/ml (unless specified) DC-SIGN-Fc (R&D Systems) or soluble polyclonal goat F(ab’)_2_ anti-human IgM (Southern Biotech) for 30 minutes on ice in 96-well (U-bottom) plates. Subsequently, 100µl containing 0.3×10^6^ cells were added into the VCAM-1-coated wells and incubated at 37°C for 30 minutes (in triplicate per condition). After 3 washes with calcium-enriched medium, adherent cells were detached using 10 mM EDTA, collected, and washed in PBS. Cells were resuspended in 200µl RPMI with 1% FBS and counted for 60 seconds using a FACS BD Canto II (BD Biosciences) at a constant medium flow rate. Data were analyzed using FlowJo software (LLC).

For adhesion assays in the presence of hIgG1-D1, 20 µg/ml DC-SIGN (equivalent to 300nM) was incubated with 500 nM hIgG1-D1 for 30 minutes on ice before cell treatment.

For adhesion assays in the presence of inhibitors, cells were treated with 0.5 µM entospletinib, 1 µM ibrutinib, 1 µM CAL-101 (Selleckchem), 50 µM CK666, 20 µM SMIFH2, or 20 µM ML141 (Sigma-Aldrich), or DMSO for 1 hour at 37°C before cell incubation with sIg ligands.

### Adhesion and Viability Assays with YK6/SIGN cells

Immortalized YK6 cells were derived from human FDCs,^25^ and were transduced with DC-SIGN to form stably expressing YK6/SIGN cells (**Supplemental Methods**). YK6/SIGN cells were irradiated and cultured with FL primary cells (at 1×10^6^ cells/ml) in the presence or absence of 300 nM hIgG1-D1 for 24 hours in 24-well plates. The fraction non-adhering to YK6/SIGN was collected by gentle pipetting, while the adherent fraction (including irradiated YK6/SIGN and infiltrated FL cells) was incubated for 15 minutes on ice in PBS before collection.

Cells from the adherent and non-adherent fractions were counted, and number of alive CD19+ cells in each condition was calculated following APC-conjugated anti-CD19 (Biolegend) staining for 30 minutes (**Table S2**).

For viability assays, anti-CD19 labelled primary FL adherent cells were subsequently stained with FITC-Annexin V and propidium iodide (PI) (Invitrogen), and the proportions of alive cells were calculated in the fraction negative for both Annexin V and PI staining.

Samples with low starting viability (<70%) or high spontaneous death during culture (more than 40% mortality difference between time 0 and 24 hours) *in vitro* were excluded from the YK6/SIGN analyses.

## RESULTS

### DC-SIGN induces low-level reorganization of sIg-Mann clusters at the cell membrane

SIg-Mann endocytosis occurs within 30 minutes of treatment at 37°C of WSU-FSCCL or primary FL cells with anti-Ig, while it does not occur with DC-SIGN (**Figure S1**). The membrane redistribution of sIg-Mann was investigated by single-molecule localization microscopy (dSTORM) following DC-SIGN or anti-Ig engagement of lymphoma cells for 10 minutes at 37°C (**Figure 1**), before complete endocytosis. DC-SIGN induced clusters with a higher number of localizations (mean 168 vs 77, p<0.0001, **Figure 1B**), and wider areas (52512 vs 34794 nm^2^, p=0.0032, **Figure 1C**) with increased density of localizations per cluster compared to untreated cells (0.0055 vs 0.0048 nm^-2^, p<0.0001, **Figure 1D**). Although the mean DC-SIGN-induced cluster area was similar to anti-Ig (52512 vs 48874 nm^2^, p=0.8153, **Figure 1B**), the number (168 vs 479, p<0.0001, **Figure 1C**) and density (0.0055 vs 0.032 nm^-2^, p<0.0001, **Figure 1D**) of localizations per cluster were significantly lower compared to anti-Ig. These data indicate that the mobility of sIg-Mann on the membrane induced by DC-SIGN is reduced compared to anti-Ig stimulation. The reduced diffusion in part explains DC-SIGN’s inability to induce sIg-Mann endocytosis (**Figure S1**),^14,15^ and low-level sIg-Mann signaling.^14,15^ However, membrane dynamics also affect integrin activation and cell adhesion. Therefore, we investigated the capacity of the DC-SIGN:sIg-Mann interaction to promote adhesion.

**Figure 1.**
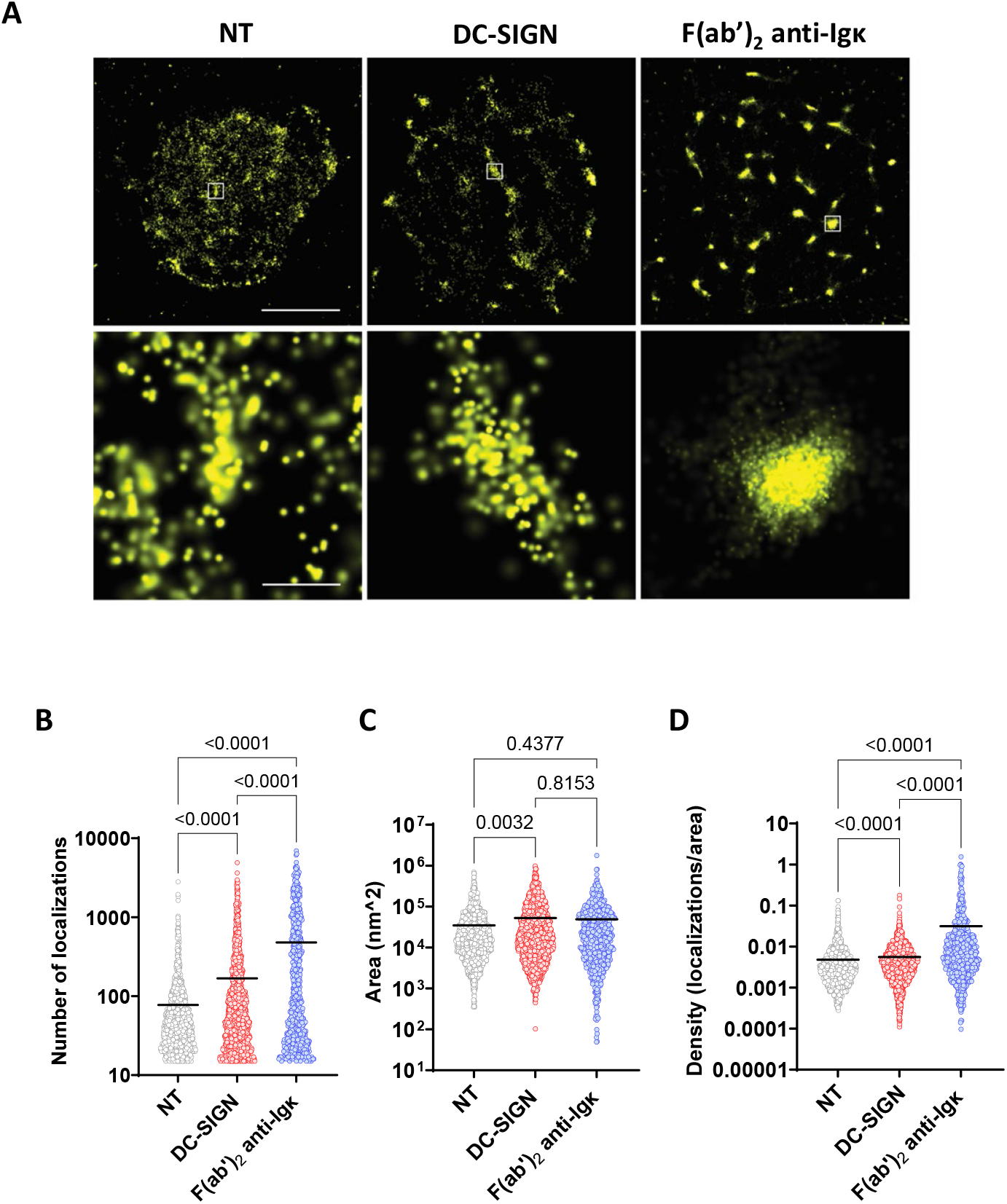
DC-SIGN induces sparse IgM clusters on the membrane of sIg-Mann+ve cells. **(A)** dSTORM reconstructed (top panels; scale bar 2 µm) and zoomed-in images (lower panels, scale bar 200 nm) of sIgM redistribution on WSU-FSCCL cells non-treated (NT), or following treatment with DC-SIGN or F(ab’)_2_ anti-Igκ. **(B-D)** Graphs showing number of localizations per cluster **(B)**,area, and **(C)** density of the clusters (calculated as the number of localizations per unit area of each cluster) **(D)** in the different conditions. Data are represented as the quantification of clusters from a total of 51 cells from 2 independent experiments. In each dot plot, the black lines indicate mean values. P-values were calculated using the Kruskal-Wallis with Dunn’s multiple comparison tests.

### DC-SIGN promotes adhesion of sIg-Mann+ve FL cells specifically to VCAM-1

VCAM-1 is highly expressed on multiple cellular and stromal elements of the lymph node, including FDCs,^26-29^ and is a candidate for adhesion. Therefore, we investigated the hypothesis that DC-SIGN interaction with sIg-Mann promotes FL cell adhesion to VCAM-1 in the sIg-Mann+ve cell line WSU-FSCCL (**Figure 2A**) and primary FL cells (**Figure 2B** and **Table S1**). Following exposure to soluble DC-SIGN, sIg-Mann-expressing cells adhered to immobilized VCAM-1 significantly more than baseline in WSU-FSCCL (p=0.0312). Pre-incubation of DC-SIGN with hIgG1-D1 antibody, which specifically blocks the DC-SIGN carbohydrate-recognition domain, completely abolished the DC-SIGN-induced adhesion. VCAM-1 adhesion assays were also performed with 3 classic FL (**Table S1**). All primary samples contained a CD20+ve/CD10+ve population with the tumor sIg that had at least 1 AGS in the CDR and bound recombinant DC-SIGN *in vitro*, consistent with the diagnosis (**Table S1** and **Figure S2)**. The analysis of the primary FL samples validated the results obtained with the cell line, by documenting that adhesion to VCAM-1 by CD20+ve/CD10+ve FL cells was significantly increased following DC-SIGN stimulation (p=0.0121) (**Figure 2B**).

**Figure 2.**
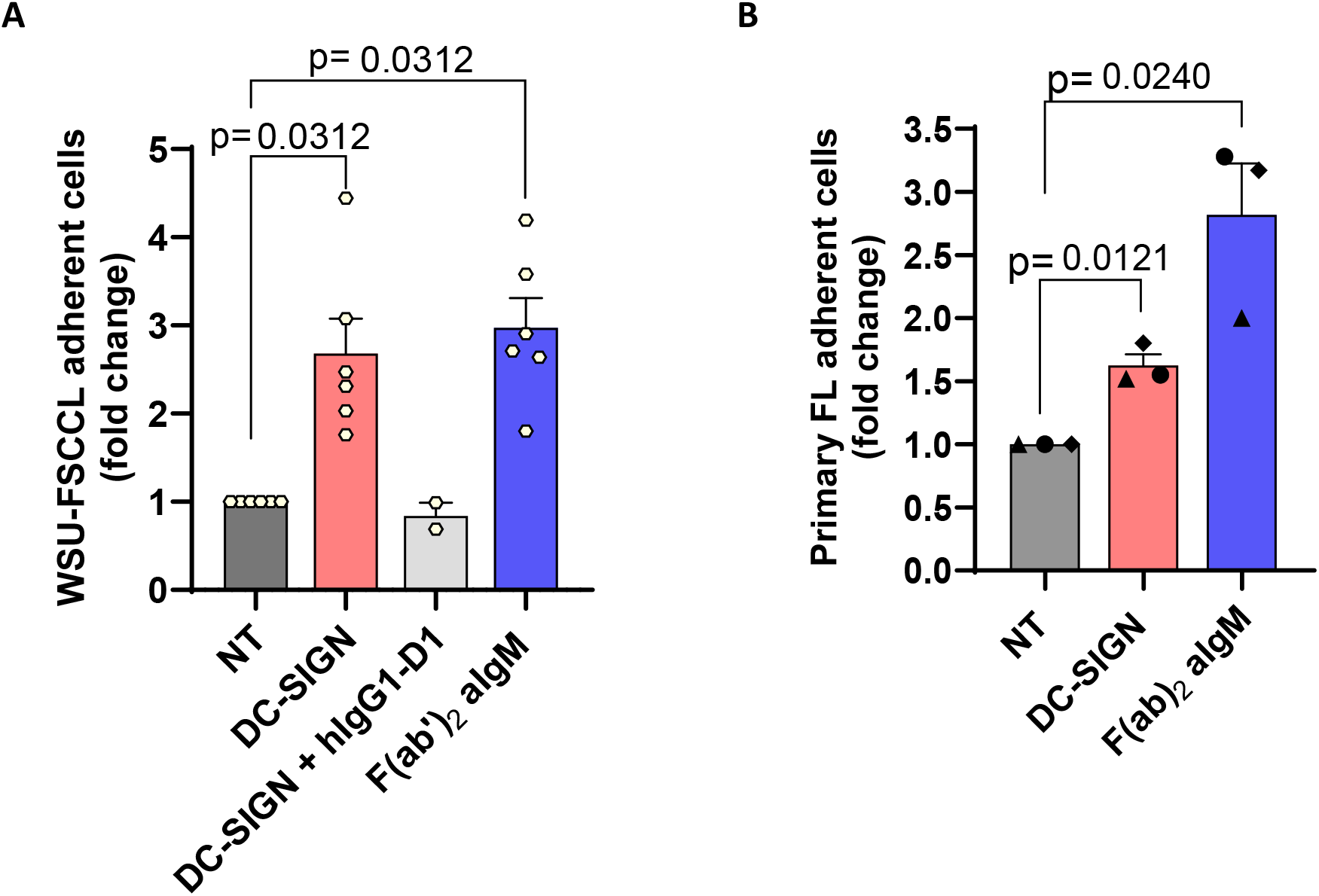
DC-SIGN induces adhesion to VCAM-1. **(A)** Adhesion of the WSU-FSCCL cell line to VCAM-1 coated on plates following treatment with F(ab’)_2_ anti-IgM, DC-SIGN alone, DC-SIGN pre-incubated with hIgG1-D1, or control (NT: non-treated). Data are represented as the mean±SEM of 6 independent experiments. Each dot represents the average of repeats in each condition per experiment. P-values were calculated using the Wilcoxon signed-rank test. **(B)** Adhesion of primary FL samples (n=3) to VCAM-1 coated on plates following treatment with F(ab’)_2_ anti-IgM, DC-SIGN, or NT control. Data are represented as mean±SEM. Each dot represents the average of repeats in each condition per experiment. P-values were calculated using the ratio-paired T-test.

### Ig-associated proximal kinases and actin polymerization control DC-SIGN-induced adhesion

The influence of signaling or actin regulators on DC-SIGN-induced adhesion was investigated with inhibitors targeting proximal Ig-associated kinases or the actin machinery. The inhibitors of SYK (entospletinib), BTK (ibrutinib), and PI3Kδ (CAL-101) all prevented the induction of adhesion by DC-SIGN (**Figure 3A-C**). CK666, which specifically inhibits the activity of the Arp2/3 complex, or SMIFH2, which inhibits formins, or CDC42 inhibitor ML141, which inhibits overall actomyosin organization, all decreased basal and DC-SIGN-induced adhesion (**Figure 3D-F**). The reduction was particularly evident with the formin inhibitor SMIFH2 (**Figure 3E**). At the concentrations used, none of the inhibitors affected cell viability for the duration of the assay (**Figure S3**). The effect of the inhibitors on DC-SIGN-induced adhesion was similar to anti-Ig, despite the inability of DC-SIGN to induce prominent sIg clustering and endocytosis.^15^ These results indicated both sIg-associated proximal kinases and actin regulators were required to induce DC-SIGN and anti-Ig-mediated adhesion, and that the DC-SIGN-mediated membrane events (including clustering) were adequate for adhesion competency of the FL cells.

**Figure 3.**
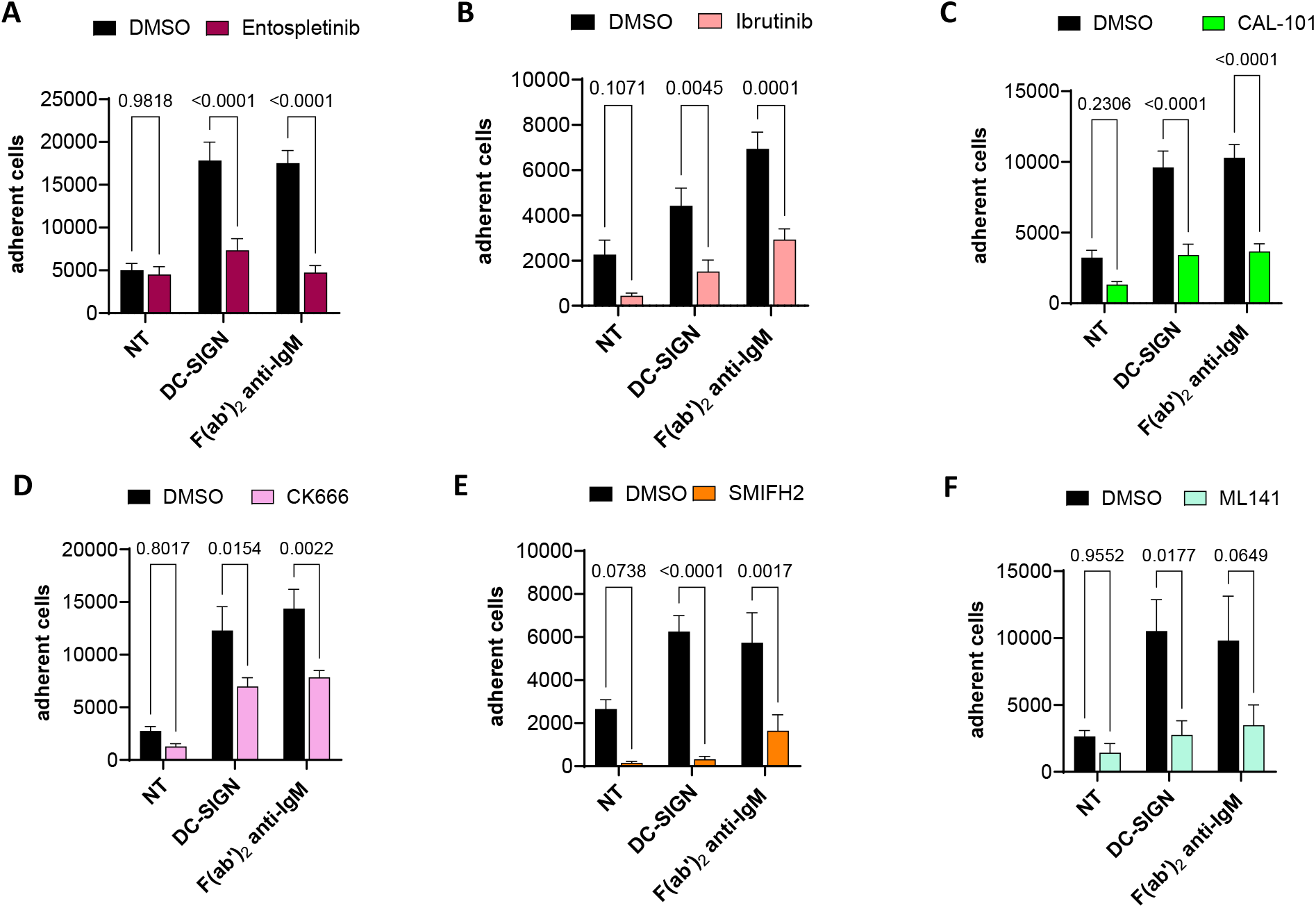
Inhibitors of early BCR signaling and actin regulators inhibit DC-SIGN-induced adhesion to VCAM-1. WSU-FSCCL cells were treated with inhibitors of molecules associated with the early BCR signaling pathway or actin polymerization, and adhesion to VCAM-1 was measured following incubation with DC-SIGN or F(ab’)_2_ anti-IgM or control (NT: non-treated). When targeting the early BCR signaling pathway, adhesion was measured following incubation with SYK inhibitor entospletinib **(A)**, BTK inhibitor ibrutinib **(B)**, or the PI3Kδ inhibitor CAL-101 **(C)**. When targeting the actin pathway, adhesion was measured following incubation with the Arp2/3 complex inhibitor CK666 **(D)**, formin inhibitor SMIFH2 **(E)**, or the CDC42 inhibitor ML141 **(F)**. Data are represented as the mean±SEM of at least 2 independent experiments. P-values were calculated using the two-way ANOVA statistical test.

### The direct interaction of sIg-Mann with DC-SIGN-expressing cells sustains adhesion of FL cells

Treatment with soluble recombinant DC-SIGN spares WSU-FSCCL (**Figure S4A**) and primary FL cells (**Figure S4B**) from apoptosis, while promoting adhesion to VCAM1. However, natural DC-SIGN is expected to operate on FL cells while expressed on macrophages and FDC, and not as a soluble molecule *in vivo*. Therefore, we assessed the direct effect of DC-SIGN expressed on a human FDC-derived cell line (YK6/SIGN) on adhesion and viability of primary FL cells, all verified for the presence of AGS in the CDR and expression of sIg-Mann by DC-SIGN binding (**Table S1**). CD19 was used to identify the FL cells (**Figure 4A**), and hIgG1-D1 antibody was used to block DC-SIGN:sIg-Mann interaction (**Figure 4B-C**).

**Figure 4.**
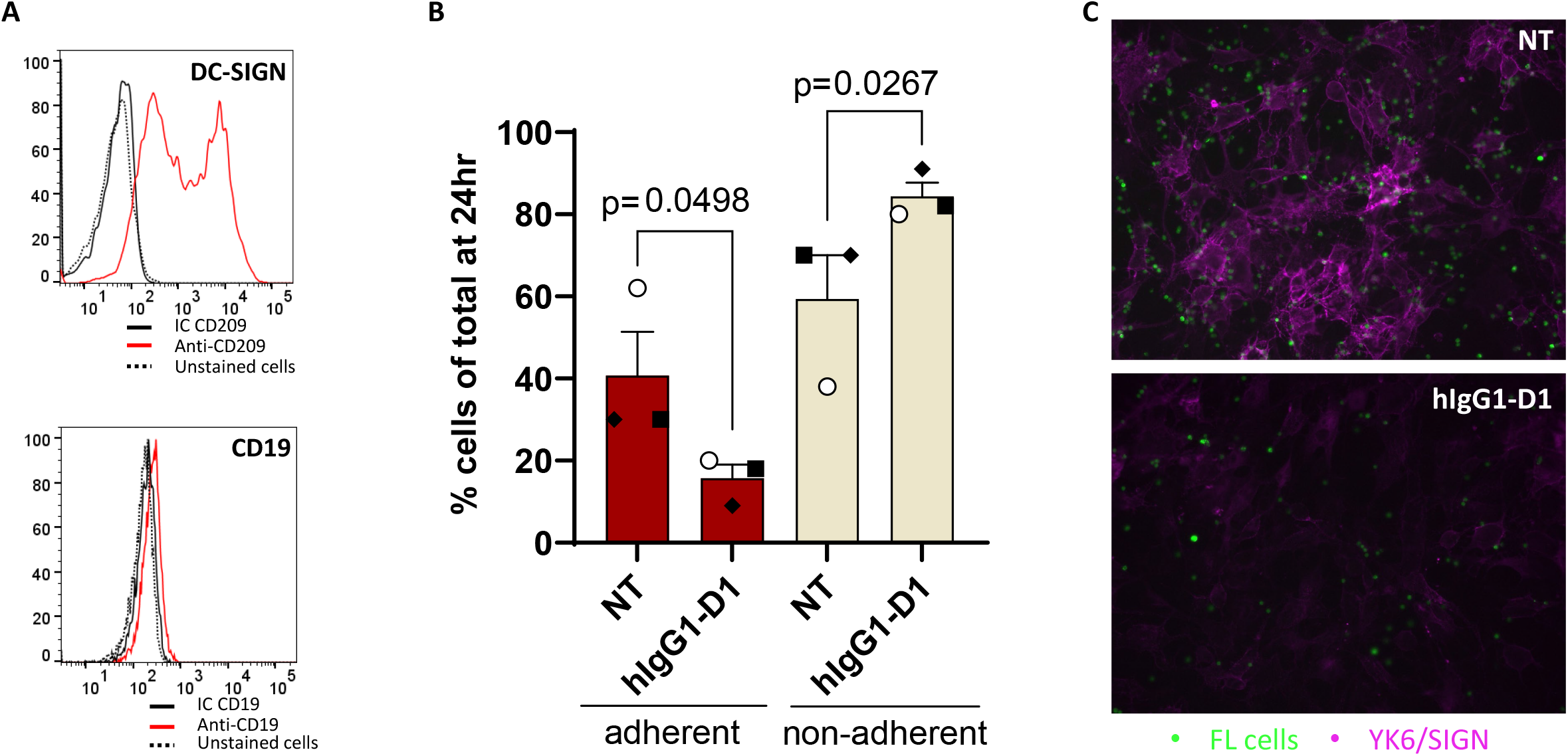
Primary FL cells adhere to co-cultured YK6/SIGN cells by direct DC-SIGN:sIg-Mann interaction. **(A)** Flow cytometry analysis of DC-SIGN (top panel) and CD19 (lower panel) expression by irradiated YK6/SIGN cells. Red lines indicate cells stained with APC-conjugated anti-CD209 or FITC-conjugated anti-CD19; black lines indicate cells stained with the relative isotype controls (IC); black dotted lines indicate autofluorescence of unstained cells. **(B)** Counts of the cells from primary FL samples (n=3) from the adherent and non-adherent fractions following 24 hours co-culture with irradiated YK6/SIGN cells in the presence of 300nM hIgG1-D1 or non-treated (NT). P-values were calculated using the ratio paired t-test. **(C)** Fluorescence microscopy images of CFSE-labeled FL cells (green, sample LY-153) in contact with YK6/SIGN cells (magenta) following 24-hour culture with YK6/SIGN in the presence/absence of hIgG1-D1. Non-adherent cells were removed by gentle pipetting. DC-SIGN on YK6/SIGN cells is shown in magenta. The dimmed staining of DC-SIGN in the hIgG1-D1-treated cells results from hIgG1-D1 partial steric interference with the detecting anti-DC-SIGN antibody (clone DCN46, BD). Magnification x20.

The effect of DC-SIGN:sIg-Mann blocking on FL cell adhesion to YK6/SIGN was assessed in FL patients’ samples (**Table S2**) by flow cytometry (**Figure 4B**) and microscopy (**Figures 4C** and **S5-S6**). Flow cytometry showed that treatment with hIgG1-D1 decreased the proportion of FL cells in contact with YK6/SIGN cells in all the primary FL samples analyzed (p=0.0498) (**Figure 4B**). Analysis of the non-adherent cell fraction was consistent with these results, since treatment with hIgG1-D1 increased the proportion of FL cells in the supernatant (p=0.0267). Microscopy analysis showed an increase of cells in suspension following treatment with hIgG1-D1 (**Figure S5**), while fluorescent imaging confirmed the decrease in the number of cells adherent to the YK6/SIGN layers (**Figure 4C and S6**). These results confirmed that blocking the DC-SIGN:sIg-Mann interaction inhibited the aggregation of primary FL cells with DC-SIGN-expressing cells.^3^

### The interaction of sIg-Mann with DC-SIGN-expressing cells sustains the survival of primary FL cells

The effect of DC-SIGN:sIg-Mann blocking on tumor viability was measured in the FL cells remaining within the YK6/SIGN layers of 8 primary FL samples by flow cytometry and fluorescence microscopy (**Table S1** and **Figure 5A-C**). The proportion of FL cells remaining alive in the non-treated (NT) condition was variable between samples. However, flow cytometric analysis revealed that treatment with hIgG1-D1 significantly increased the proportion of non-viable cells compared to untreated cells in all 8 samples (p=0.0078, **Figure 5B**). Consistently, fluorescent imaging confirmed that the majority of the residual cells remaining within the YK6/SIGN layers were apoptotic following hIgG1-D1 treatment (**Figure 5C**).

**Figure 5.**
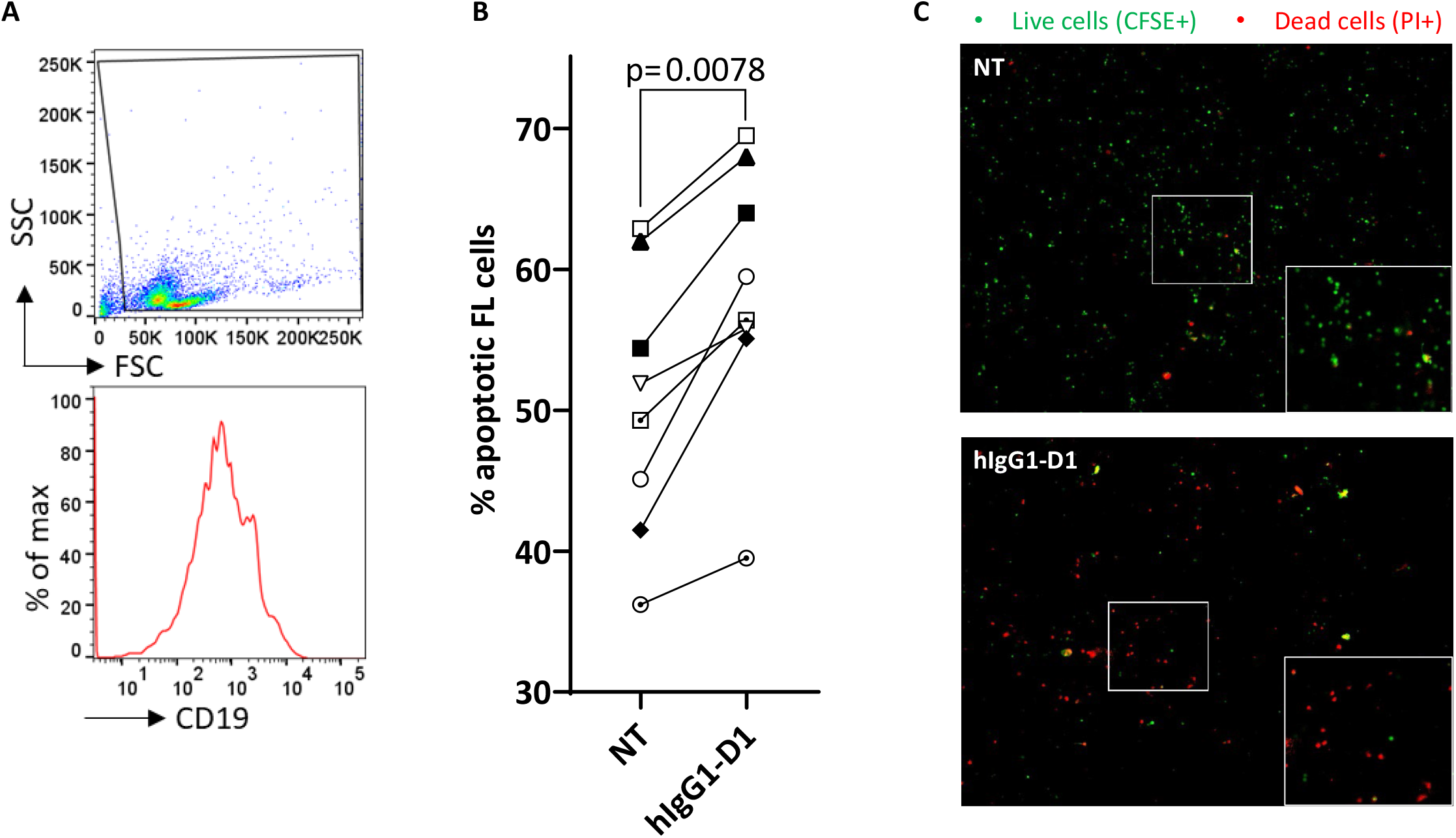
The viability of primary FL cells co-cultured with YK6/SIGN cells is inhibited by blocking the DC-SIGN:sIg-Mann interaction. **(A)** Gating strategy to determine cell viability in CD19+ve primary FL cells in contact with irradiated YK6/SIGN cells (19-TB0260 is shown as an example): FL cells treated with hIgG1-D1 (300nM) or non-treated in the co-culture were identified using CD19 marker and viability was determined by AnnexinV/PI staining of the CD19+ve adherent fraction by flow cytometry. **(B)** Primary FL samples (n=8) cultured with YK6/SIGN cells for 24 hours in the presence of hIgG1-D1 or non-treated (NT). The graph shows the percentage of apoptotic cells (100 – [% Annexin V-ve/PI-ve] FL cells in the CD19+ gate) in the adherent fraction at 24 hours. The p-value was calculated using the Wilcoxon signed-rank test. **(C)** CSFE-labeled primary FL cells (LY-153) were cultured with YK6/SIGN cells for 24 hours in the presence of hIgG1-D1 or no treatment (NT). Cells were visualized by fluorescence microscopy following PI staining. The images show viable FL cells in green (CFSE+ve) and dead cells in red (PI+ve), following treatment with hIgG1-D1 (bottom image) or no treatment (NT, top panel). Magnification x20. The white squares define zoomed-in areas.

These data indicated that blocking the interaction of sIg-Mann with DC-SIGN expressed on the FDC-derived cells critically affected both adhesion and survival of primary FL cells.

## DISCUSSION

These data provide new fundamental information on how the unique and tumor-specific interaction of sIg-Mann with DC-SIGN supports FL cell retention and survival in the tumor lymph node.

Occupation of the antigen-binding site by oligomannose-type glycans is a tumor-specific feature of FL.^22^ The microanatomical features indicate that the FL cells preferentially reside in the lymph node, where macrophages and FDCs expressing DC-SIGN are present in variable amounts.^7^

In infection, the key role of FDCs is to present complement-opsonized antigens to sIg on germinal center B cells, which have transited to the light zone for selection. Antigen-specific B cells receive survival signals when they bind to the opsonized antigen in the presence of T-cell help. However, the interaction modalities of FL cells with FDCs in the absence of antigen appear different. The occupation of the antigen-binding site by oligomannose-type glycans prevents binding to conventional antigens,^8^ and mutations of *CREBBP* or *EZH2* affect T-cell help by reducing MHC-II on the FL cell surface and by diverting B-cell dependency in the light zone from T-follicular helper cells to FDCs.^30,31^ In our co-culture model *in vitro*, we find that the primary FL cells adhere to DC-SIGN-expressing FDC-derived cells, and the specific DC-SIGN:sIg-Mann interaction confers a survival advantage. If the interaction is inhibited by an agent specifically blocking the carbohydrate-recognition domain of DC-SIGN, FL cells lose adherence and/or the tumor cells remaining within the FDC-derived cell layers die. Therefore, our data indicate that the sIg-Mann on FL cells will interact with DC-SIGN on FDC to receive specific direct signals for cell survival.

The tumor-specific, antigen-independent, DC-SIGN:sIg-Mann interaction contrasts with conventional antigen:sIg protein ligation. The latter may need to be prevented in FL to avoid sIg endocytosis and cell death, which naturally occurs in the absence of T-cell help.^9,10^ In our experiments, soluble DC-SIGN does not trigger cell death. Remarkably, DC-SIGN appears to mimic the interaction of monovalent ligands in their inability to promote sIg endocytosis, and *in vitro* pre-exposure to DC-SIGN protects the sIg from further stimulation, keeping the cells under a persistent low-level lymphoma-promoting signal.^14,15^ Low-level sIg-mediated signals are essential for B-cell survival,^1,20^ but still require membrane skeleton modifications to control adequate signaling through the B-cell receptor.^32^ We find that DC-SIGN is able to modify the redistribution of sIg at the plasma membrane, in a different manner from F(ab’)^2^ anti-Ig, further explaining the absence of endocytosis, the low-level sIg stimulation, while the signals are adequate to sustain adhesion to VCAM-1.

We had already observed that the specific DC-SIGN:sIg-Mann interaction promotes the formation of sIg-Mann+ve cell clusters around DC-SIGN-expressing cells.^3^ However, VCAM-1 is highly expressed on FDC and macrophages,^26-29^ and the mechanisms by which sIg engagement mediates adhesion have been well studied in lymphoma cell lines.^33^ This study confirms a direct effect of DC-SIGN on the adhesion of primary FL cells to an FDC-like YK6/SIGN cell line by ligation to sIg-Mann. We provide an additional way by which DC-SIGN determines FL cell retention in the lymph node via VCAM-1. Although the signal strength of DC-SIGN is different from anti-Ig or conventional antigens,^2,34^ the pathway involved requires BTK and PI3K kinases, as expected.^33,35,36^ Our data already indicate that DC-SIGN:sIg-Mann interruption is sufficient to deprive primary FL cells of pro-survival signals and accelerate their death *in vitro*. Therefore, a therapeutic approach interrupting DC-SIGN:sIg-Mann interaction may have an anti-tumor effect in patients.

In conclusion, this study describes key functional consequences of DC-SIGN interaction with FL cells. The variable amounts of DC-SIGN on macrophages and FDCs promote local retention at tissue sites and survival of FL cells. Blocking DC-SIGN interaction with sIg-Mann inhibits these functions and may be a new therapeutic approach against FL in patients.

## ACKNOWLEDGMENTS

We are very grateful to Freda Stevenson, Professor Emeritus at the University of Southampton, for her continued mentorship and support over the years. We thank Dr Kathy Potter and the Cancer Sciences Tissue Bank for the provision of primary cell suspension for functional assays. This work was funded by the Eyles Cancer Immunology post-doctoral fellowship and the Eyles PhD fellowship, Cancer Research UK BTERP (C36811/A29101), Cancer Research UK Accelerator ECRIN-M3 (C42023/A29370), and CRUK programme grant (C2750/A23669). G.C. was funded by the Leukaemia UK John Goldman Fellowship.

## AUTHORSHIP CONTRIBUTIONS

G.C. performed research, analyzed and interpreted data, and wrote the manuscript. D.T., L.R, E.S., S.J., and S.L. performed experiments. P.J.D. generated primary reagents. M.D.R. and D.H. generated the YK6/DC-SIGN cells. P.R. and R.B. diagnosed, selected, and provided primary FL samples. M.V. provided the hIgG1-D1 antibody. F.F. designed the study, supervised research, interpreted data and wrote the manuscript. All authors reviewed and approved the manuscript.

