## Supplemental material for "DC-SIGN Binding to the Surface Ig Oligomannose-type Glycans Promotes Follicular Lymphoma Cell Adhesion and Survival"

### SUPPLEMENTAL METHODS

*Patients and cell lines*

Primary sample material was obtained from patients with FL at the Department of Pathology, University of Rochester Medical Center, and at the University of Southampton in the Mature Lymphoid Malignancies Observational Study (NIHR/UKCRN Portfolio ID: 31076). For phenotyping, immunoglobulin gene analysis and functional assays, FL cell suspensions from fresh tumor lymph nodes cryopreserved in fetal bovine serum (FBS) + 10% dimethyl sulfoxide were used.

WSU-FSCCL cell line was obtained from DSMZ and cultured as previously described.[1]

FDC-like YK6 cells were previously immortalized through retroviral transduction with pBABE_TERT.Hygro, P53DD_Thy1.1 and CDK4_R24C_Thy1.1.[2] The CDS encoding DC-SIGN (24-1238, (NM_021155, Genescript) was cloned into a pBMN-IRES-LyT2 retroviral vector and the immortalized YK6 cells were further transduced to generate DC-SIGN expressing cells (YK6/SIGN). Expression of DC-SIGN was confirmed by flow cytometry. Cells were grown in RPMI supplemented with 10% FBS.

*Phenotyping and DC-SIGN binding assay*

Before each assay, viable cryopreserved cells were purified post-thawing by Lymphoprep (Stemcell Technologies) gradient centrifugation. Phenotype and DC-SIGN binding of primary FL specimens were performed by flow cytometry, as previously described.[1] Briefly, cells were stained on ice for 30 minutes with APC-conjugated anti-CD10 (Biolegend), PerCP-Cy5.5-conjugated anti-CD20 (Biolegend), FITC-conjugated $F(ab')_2$ anti-IgG (Dako), PE-conjugated $F(ab')_2$ anti-IgM (Dako), or respective controls. DC-SIGN binding was performed by treating cells with 20µg/ml soluble recombinant DC-SIGN-Fc or DC-SIGN-HA (R&D Systems) in calcium-enriched medium before incubation with FITC-conjugated anti-human Fc or AF488-conjugated anti-HA (Biolegend), respectively. Data were acquired using a FACS BDCanto II and analyzed with FlowJo software. Both phenotype and DC-SIGN binding were analyzed in the CD10+ve/CD20+ve tumor population.

Irradiated YK6/SIGN cells were incubated for 10 minutes on ice before detachment by cell scraping. Cells were then stained for 30 minutes on ice with APC-conjugated anti-CD209 (Biolegend) or FITC-conjugated anti-CD19 (Biolegend) or the respective controls. Data were acquired using a FACS BDCanto II and analyzed with FlowJo software.

*Immunoglobulin gene analysis*

The *IG* heavy chain transcripts from primary FL specimens were obtained by sequencing as previously described.[1, 3, 4] The IMGT database and the IMGT/V-QUEST tool were used to determine homology to the closest germline sequence. The deduced amino acid sequences were analyzed for the presence of AGS (NxS/T; x ≠ P), and site location was determined using the IMGT numbering system.

*Endocytosis assay*

WSU-FSCCL or primary FL cells ($0.3x10^6$ cells/well in 100µl calcium-enriched medium) were treated with 20µg/ml soluble DC-SIGN-Fc, soluble polyclonal goat $F(ab')_2$ anti-human Igκ (Southern Biotech) or non-treated for 30 minutes on ice before incubation at 37°C for 10 or 30 minutes. Cells were then washed in cold medium and stained on ice for 30 minutes with PE-conjugated $F(ab')_2$ anti-IgM (Dako) or FITC-conjugated anti-IgG (Dako).

*Immunoblotting*

WSU-FSCCL cells were treated for 1 hour with inhibitors at 37°C, plated into U-bottom 96-well plates and incubated 20µg/ml soluble polyclonal goat $F(ab')_2$ anti-human IgM or left untreated for 30 minutes on ice in RPMI-1640 with 1% FBS and 1 mM calcium. Immunoblot analysis was then performed as previously described.[1] The following primary antibodies were used: anti-phosphoAKT (Ser473), anti-AKT, anti-phosphoBTK (Tyr223), anti-BTK, anti-phosphoSYK (Tyr525/526), anti-SYK (all from Cell Signaling Technology), and anti-GAPDH (6C5) (Ambion). After washing, membranes were stained with horseradish peroxidase (HRP)-conjugated secondary antibodies (Dako), and images were acquired using the Amersham ImageQuant800 (GE Life Sciences).

*Viability assays*

WSU-FSCCL cells ($0.125x10^6$ per well) were incubated in 250 µl calcium-enriched medium with 20µg/ml soluble DC-SIGN-Fc or no treatment (NT) in 96-well flat-bottom plates at 4°C for 30 minutes, and then at 37°C for 6 hours. Primary FL samples ($0.25x10^6$ cells per well) were incubated in 100 µl calcium-enriched medium with 20µg/ml soluble DC-SIGN-Fc or no treatment (NT) in 96-well flat-bottom plates at 37°C for 24 hours. Cells were washed and stained with FITC-Annexin V and propidium iodide (PI) (Invitrogen) in 1X Annexin binding buffer. Viability was measured by flow cytometry using a FACS BDCanto II.

To assess viability of cells undergoing adhesion assays, cells were incubated in calcium-enriched medium with 1% FBS and inhibitors for 1 hour at 37°C, then on ice for 30 minutes, and 30 minutes at 37°C prior to Annexin V/PI staining.

*Fluorescence microscopy*

FL cells were labeled with CFSE (10 minutes at 37°C in PBS) and washed 3 times in RPMI-1640 with 10% FBS before co-culture with irradiated YK6/SIGN cells ($0.5x10^6$ FL cells in 500µl per well) in presence/absence of 300nM hIgG1-D1. After 24-hour incubation, non-adherent cells were removed by gentle pipetting and cells were fixed with 0.5% PFA for 10 minutes at room temperature. Cells were then washed and blocked for 30 minutes at room temperature, before incubation with 5µg/ml anti-DC-SIGN antibody (clone DCN46, BD Biosciences) for 1 hour at room temperature. Following incubation, cells were washed and stained with AF568-conjugated anti-mouse secondary antibody (1:250, Invitrogen) for 45 minutes at room temperature. Cells were then washed and imaged with an EVOS M5000 microscope. Images were analyzed with the ImageJ software.

To assess viability of cells by fluorescence microscopy, CFSE-labeled cells were cultured with irradiated YK6/SIGN cells as described above. Following 24-hour incubation, non-adherent cells were removed by gentle pipetting while adherent cells were stained with PI (1:500 in PBS) for 5 minutes at room temperature. Cells were washed 3 times in PBS before imaging with an EVOS M5000 microscope.

*Statistics*

Analyses were performed in Prism 9.4.1 (GraphPad Software). Continuous variables were compared using the ratio paired t-test or Wilcoxon signed rank test, as appropriate. Changes across different levels of treatment and conditions were analyzed using a two-way ANOVA. Categorical variables were compared using the Chi-square statistical test.

### SUPPLEMENTAL TABLE LEGENDS

**Table S1. Characteristics of follicular lymphoma patients used in the study.**

**Table S2. Analysis of adhesion of primary FL cells to YK6/SIGN cells following co-culture for 24 hours in the absence (NT) or presence of hIgG1-D1 antibody.**

### SUPPLEMENTAL FIGURES LEGENDS

**Figure S1. Endocytosis of sIg-Mann occurs following anti-Ig but not DC-SIGN treatment.** Cells were treated with anti-Ig or DC-SIGN for 30 minutes on ice before incubation for 10 or 30 minutes at 37°C. **(A)** WSU-FSCCL cells (IgM+ve) were treated with F(ab')$_2$ anti-Igκ or DC-SIGN and sIg was detected by staining with PE-conjugated anti-IgM or control. **(B)** Primary FL cells from sample LY-153 (IgG+ve) were treated with F(ab')$_2$ anti-Igκ or DC-SIGN and sIg was detected by staining with FITC-conjugated anti-IgG or control. In A and B, red lines indicate cells treated with soluble DC-SIGN and stained with either PE-conjugated anti-IgM or FITC-conjugated anti-IgG; blue lines indicate cells treated with anti-Ig and stained with either PE-conjugated anti-IgM or FITC-conjugated anti-IgG; grey filled histograms indicate cells non treated with sIg ligands and stained with either PE-conjugated anti-IgM or FITC-conjugated anti-IgG; black lines indicate cells non treated with sIg ligands and stained with either PE-conjugated control or FITC-conjugated control; black dotted lines indicate the autofluorescence of unstained cells. Colored numbers indicate mean fluorescent intensity for each condition.

**Figure S2. *In vitro* characterization of the FL specimens used in the study.** Tumor cells of the 9 FL samples used in the study (Table S1) were characterized by sequencing of the IGHV-D-J rearrangement, DC-SIGN binding to sIg-Mann and sIg phenotyping. Two representative samples are shown. IGHV-D-J deduced amino acid sequence **(A)**, gating strategy of the CD10+/CD20+ tumor population **(B)**, DC-SIGN binding **(C)** and sIg expression **(D)** of sample 13-TB0582 are shown in the upper panels. The deduced amino acid sequence **(E)**, gating strategy of the CD10+/CD20+ tumor population **(F)**, DC-SIGN binding **(G)** and sIg expression **(H)** of sample 19-TB0260 are shown in the lower panels. In panels C and G, red lines indicate tumor cells stained with soluble DC-SIGN-Fc followed by FITC-conjugated anti-Fc; black lines indicate tumor cells not treated with soluble recombinant DC-SIGN-Fc and then stained with FITC-conjugated anti-Fc (NT); black dotted lines indicate the autofluorescence of unstained cells. In panels D and H, red lines indicate cells stained with either PE-conjugated anti-IgM or FITC-conjugated anti-IgG; black lines indicate cells stained with the respective polyclonal controls (ctrl); black dotted lines indicate the autofluorescence of unstained cells. IC: isotype control.

**Figure S3. Cell viability during the adhesion assays in the presence of inhibitors. (A)** Flow cytometry plots showing the viability of WSU-FSCCL following incubation with inhibitors targeting the early BCR signaling pathway (entospletinib, ibrutinib, and CAL-101) or the actin/cytoskeletal pathways (CK666, SMIFH2, and ML141). **(B)** Immunoblotting showing inhibition of protein phosphorylation following treatment with kinase inhibitors and subsequent sIg stimulation with F(ab')$_2$ anti-IgM (aIgM) in WSU-FSCCL cells. GAPDH was used as a loading control. Superscript numbers indicate proteins imaged in the same gel membrane.

**Figure S4. Soluble DC-SIGN spares sIg-Mann+ve cells from apoptosis. (A)** Representative histograms of WSU-FSCCL cells viability following 6-hour DC-SIGN treatment, as measured by Annexin V staining. The percentage of viable cells is shown in the top left corner of each plot. The graph chart shows the percentage of alive WSU-FSCCL cells (Annexin V-negative percentage cells) as mean ± SD of 4 independent experiments. **(B)** Histograms showing viability of two primary FL samples (LY-221 in top panels, and 11-TB0536 in bottom panels) following 24-hour DC-SIGN treatment, as measured by Annexin V staining. The percentage of viable cells is shown in the top left corner of each plot. The graph chart shows the percentage of alive FL cells (Annexin V-negative percentage cells) as mean ± SD of 2 independent experiments.

**Figure S5. Increased number of non-adherent cells in the supernatant following anti-DC-SIGN treatment.** Light microscopy images of the non-adherent fraction of unstained FL cells (sample LY-153) on the YK6/SIGN layer following 24-hour incubation with hIgG1-D1 or non-treated (NT). The amount of FL cells (light gray dots) in suspension is higher in the presence of the hIgG1-D1 antibody compared to NT. The YK6/SIGN cells (dark gray cell layer, red arrows) can be seen in the NT condition, admixed with adhering FL cells, while they are not visible when the co-culture is in the presence of hIgG1-D1, because covered by suspended cells remaining in the supernatant. Images were taken with an EVOS M5000 microscope, magnification x10.

**Figure S6. Adhesion of FL cells to YK6/SIGN is decreased by anti-DC-SIGN treatment.** Fluorescence microscopy as per Figure 4C of 2 additional field images of CFSE-labeled FL cells (green, sample LY-153) in contact with YK6/SIGN cells (magenta) following 24-hour culture with YK6/SIGN in the presence/absence of hIgG1-D1. Non-adherent cells were removed by gentle pipetting. DC-SIGN on YK6/SIGN cells is shown in magenta. The dimmed staining of DC-SIGN in the hIgG1-D1-treated cells results from hIgG1-D1 partial steric interference with the detecting anti-DC-SIGN antibody (clone DCN46, BD). Magnification x20.
