## Supplemental Figures for "DC-SIGN Binding to the Surface Ig Oligomannose-type Glycans Promotes Follicular Lymphoma Cell Adhesion and Survival"

#### Figure S1

**A**

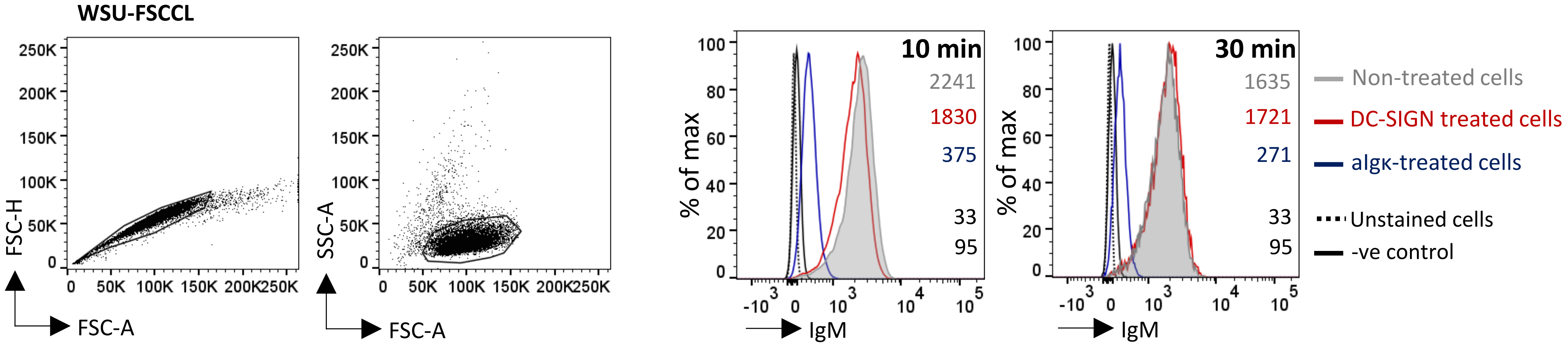

WSU-FSCCL
FSC-H
FSC-A
SSC-A
FSC-A
10 min
2241
1830
375
33
95
% of max
IgM
30 min
1635
1721
271
33
95
% of max
IgM
Non-treated cells
DC-SIGN treated cells
aIgκ-treated cells
Unstained cells
-ve control


**B**

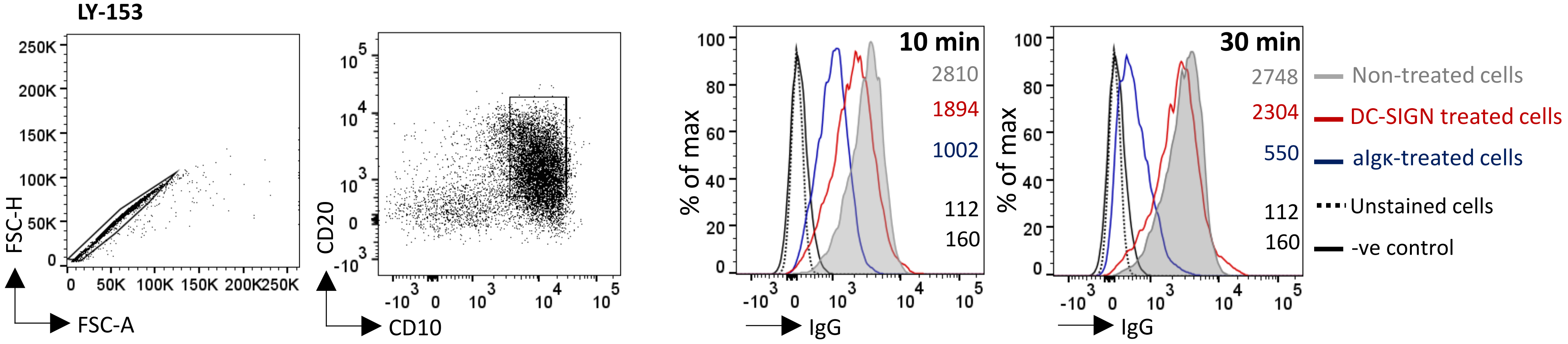

LY-153
FSC-H
FSC-A
CD20
CD10
10 min
2810
1894
1002
112
160
% of max
IgG
30 min
2748
2304
550
112
160
% of max
IgG
Non-treated cells
DC-SIGN treated cells
aIgκ-treated cells
Unstained cells
-ve control

### Figure S2

**A**

```
                                          FR1                  CDR1          FR2              CDR2              FR3                               CDR3
                                         (1-26)               (27-38)       (39-55)          (56-65)          (66-104)                          (105-117)
                               __________________________ ____________ _________________ __________ _______________________________________ __________________
                               1        10        20        30          40        50          60      70        80        90        100          111   112
                               .........|.........|...... ...|........ .|.........|..... ....|..... ....|.........|.........|.........|.... ......|     |.....
M95114 Homsap IGHV4-59*03 F    QVQLQESGP.GLVKPSETLSLTCTVS GGSI....SSYY WSWIRQPPGKGLEWIGY IYYS...GST NYNPSLK.SRVTISVDTSKNQFSLKLSSVTAADTAVYYC A          YYYYGMDV W IGHJ6*02
13-TB0582                      ---------.----V------M---- ----....-D-H ------S---S---M-N -S--...-D- -------.------LE-----V-------------I--- -RHLRGITGGS-D----- W
```

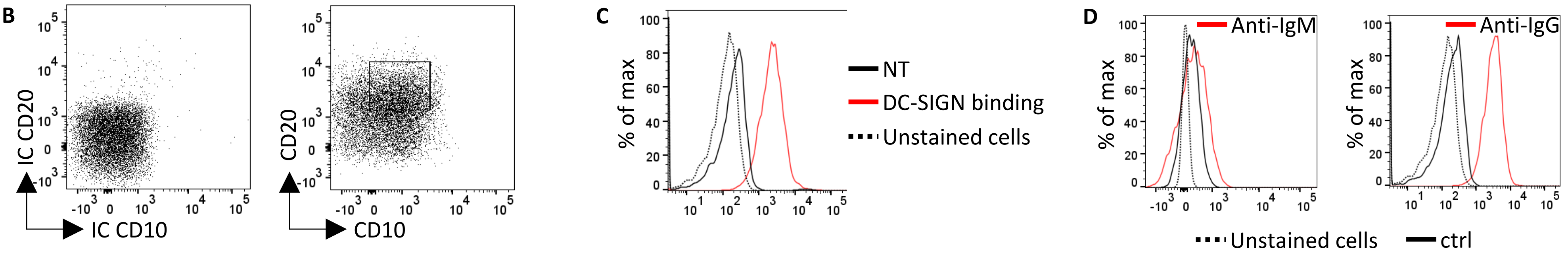


**E**

```
                                          FR1                  CDR1          FR2              CDR2              FR3                               CDR3
                                         (1-26)               (27-38)       (39-55)          (56-65)          (66-104)                          (105-117)
                               __________________________ ____________ _________________ __________ _______________________________________ ________________
                               1        10        20        30          40        50          60      70        80        90        100          111 112
                               .........|.........|...... ...|........ .|.........|..... ....|..... ....|.........|.........|.........|.... ......|   |.....
M99660 Homsap IGHV3-23*01 F    EVQLLESGG.GLVQPGGSLRLSCAAS GFTF....SSYA MSWVRQAPGKGLEWVSA ISGS..GGST YYADSVK.GRFTISRDNSKNTLYLQMNSLRAEDTAVYYC AK          YFDY W IGHJ4*02
19-TB0260                      ---------.---------------- ----....---- ----------------V ----..--T- -------.---N-----T--------------------- --NGSRGYYDSSGYY- W
```

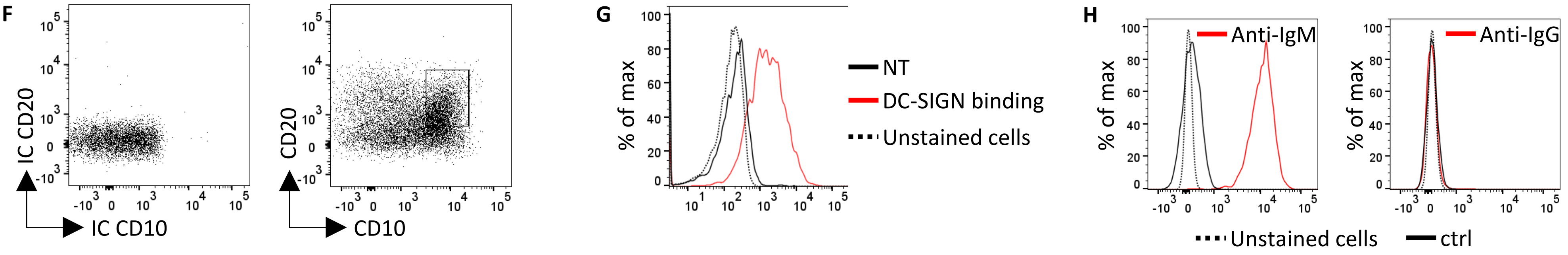

### Figure S3

A

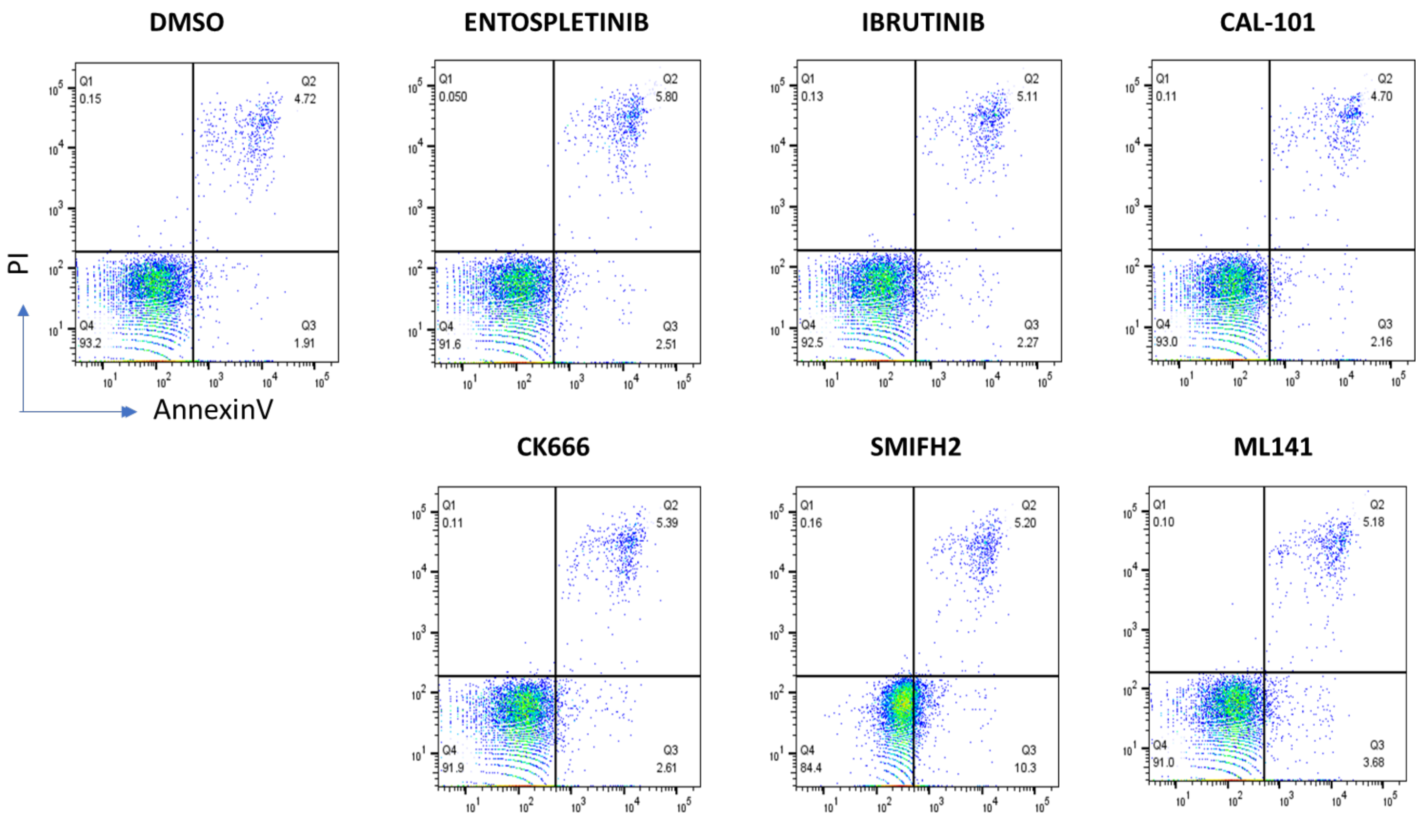
DMSO
ENTOSPLETINIB
IBRUTINIB
CAL-101
CK666
SMIFH2
ML141
PI
AnnexinV
Q1 0.15
Q2 4.72
Q4 93.2
Q3 1.91
Q1 0.050
Q2 5.80
Q4 91.6
Q3 2.51
Q1 0.13
Q2 5.11
Q4 92.5
Q3 2.27
Q1 0.11
Q2 4.70
Q4 93.0
Q3 2.16
Q1 0.11
Q2 5.39
Q4 91.9
Q3 2.61
Q1 0.16
Q2 5.20
Q4 84.4
Q3 10.3
Q1 0.10
Q2 5.18
Q4 91.0
Q3 3.68

B

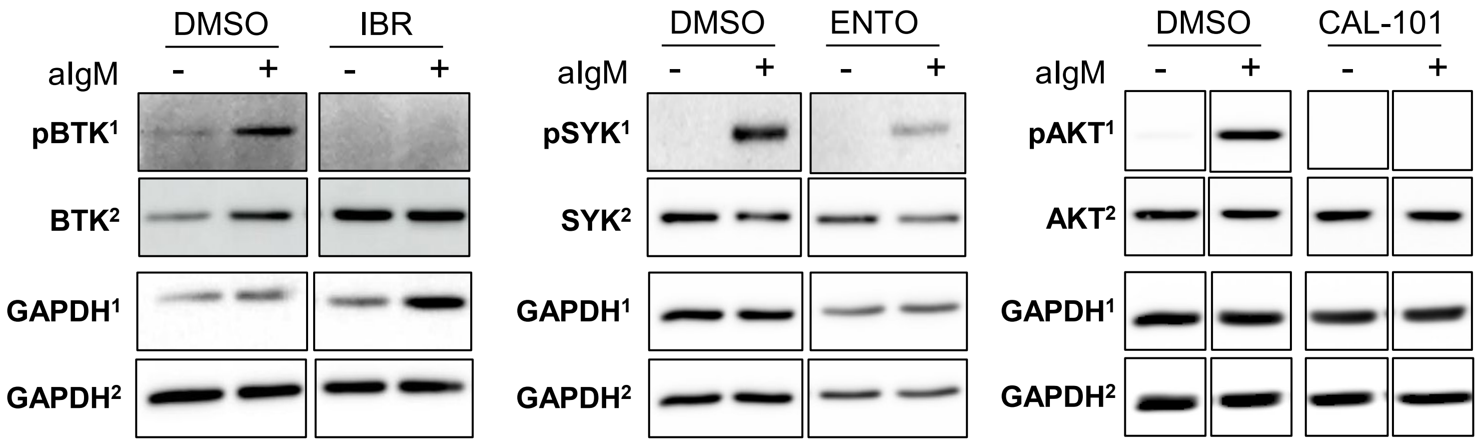
DMSO
IBR
aIgM
- + - +
pBTK1
BTK2
GAPDH1
GAPDH2
DMSO
ENTO
aIgM
- + - +
pSYK1
SYK2
GAPDH1
GAPDH2
DMSO
CAL-101
aIgM
- + - +
pAKT1
AKT2
GAPDH1
GAPDH2

**Figure S4**

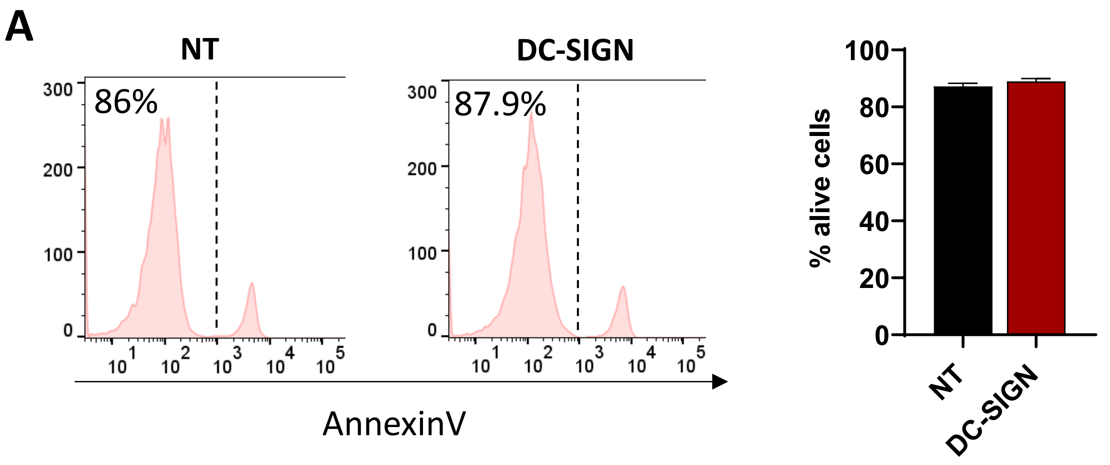
A
NT
DC-SIGN
86%
87.9%
AnnexinV
% alive cells
NT
DC-SIGN

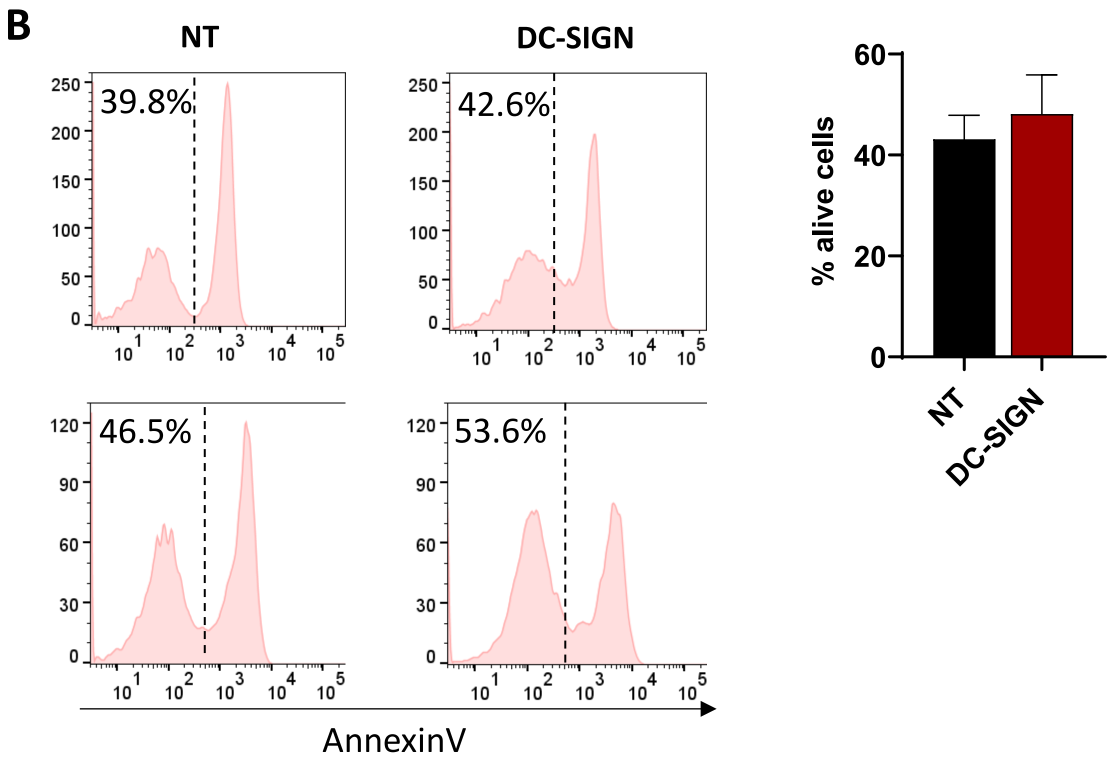
B
NT
DC-SIGN
39.8%
42.6%
46.5%
53.6%
AnnexinV
% alive cells
NT
DC-SIGN

**Figure S5**

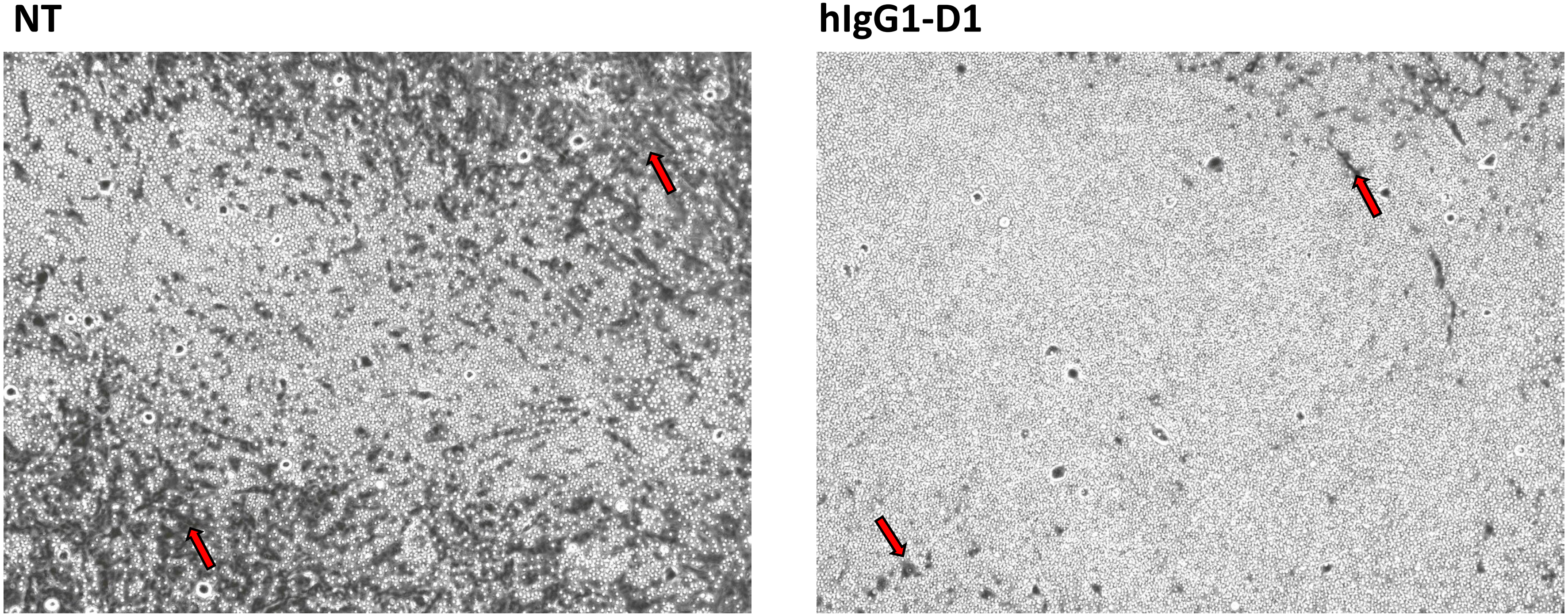

**Figure S6**

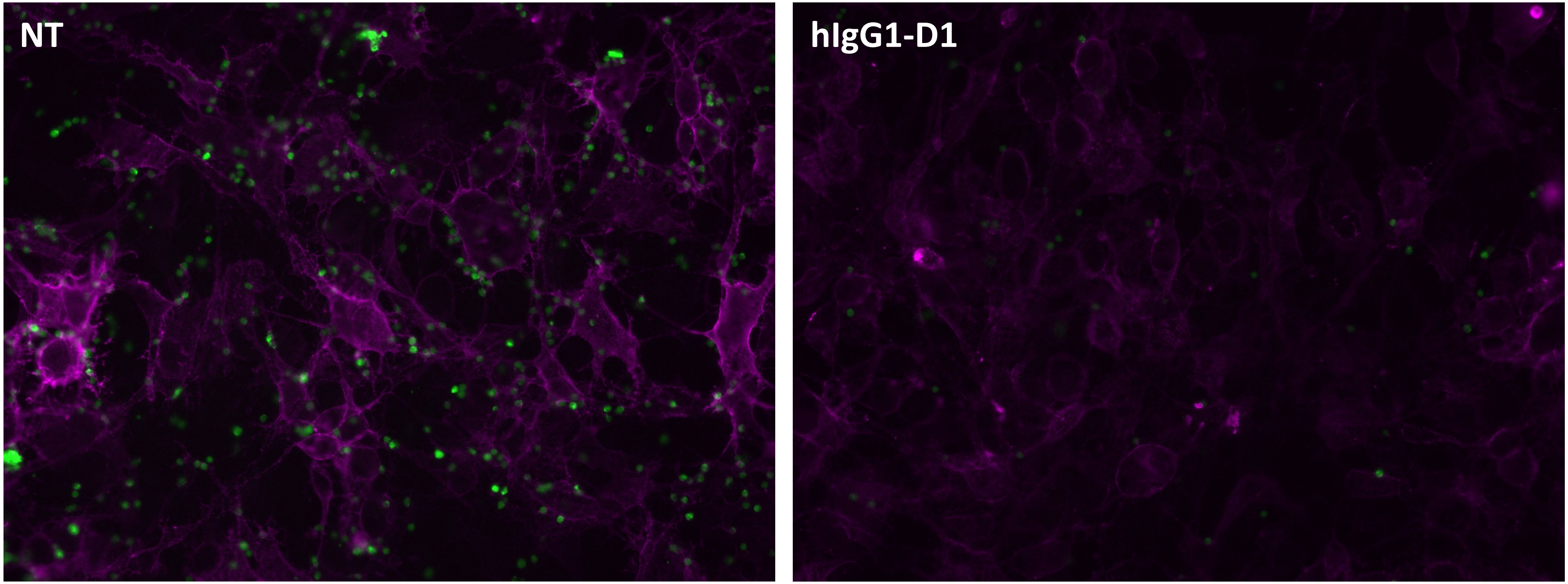


- FL cells
- YK6/SIGN
